# Geometric morphometric analysis reveals cranial shape divergence and asymmetry in extinct and extant species of big cats (Carnivora: Felidae)

**DOI:** 10.1101/2022.05.20.492788

**Authors:** Aidan Noga, William Anyonge, Amanda K. Powers

## Abstract

Felidae, a family of the order Carnivora, includes extinct and extant species of cats spread across a wide ecological and geographical landscape. Cats are well-suited for predation due to various physical and behavioral characteristics, such as optimized limb length, skull shape, as well as enhanced hearing and vision. Morphological changes across Felidae species, particularly changes in skull shape, are likely explained by differences in predatory and feeding behaviors. Toward that end, cranial shape was analyzed across six different extant and extinct Felidae species using two-dimensional geometric morphometrics. From the lateral cranial view, we discovered that the cheetah (*Acinonyx jubatus*) and the North American Sabretooth (*Smilodon*) had the most significant shape divergence, specifically at the frontal bone and post orbital regions of the skull. Specifically, we found that the Sabretooth had a significantly shorter coronoid process compared to other Felids. We also observed a significant difference in post orbital shape in the cheetah dorsal cranium. Interestingly, we found that both the cheetah and the extinct North American Lion demonstrate significant shape asymmetry in the postorbital region from a ventral view of the skull. Shape divergence and asymmetry in select Felid skulls may arise from decreased genetic diversity. Taken together, we reasoned that morphological changes in skull shape likely evolved to support differences in predatory behavior across Felidae.

## Introduction

Felidae, the family within the order Carnivora [1] includes cat species that occupy a diverse range of habitats in all parts of the world, apart from Antarctica and Australia [2]. Originating during the Eocene Epoch [3], the relatively recent emergence of this family (28.5-35 Mya) makes them an interesting evolutionary model for studying morphological divergence [4-5].

The Felidae family is comprised of two distinct evolutionary lineages, including extant modern cats and extinct sabretooth cats [6]. Some of the earliest felids to emerge were sabretooth cats, such as members of the *Smilodon* genus that inhabited North and South America during the Pleistocene Epoch [1]. Across extant species of felids, eight clades have been established using molecular evidence. These clades include the ocelot lineage, domestic cat lineage, *Panthera* genus, puma group, *Lynx* genus, Asian leopard cat group, caracal group, and the bay cat group [5,7]. The earliest split among these living felids occurred roughly 10.8 Mya with the emergence of *Panthera* genus [5] during the Miocene and Pliocene Epoch [8]. Notable members of this species include the African lion, leopard, tiger, snow leopard, and extinct North American (NA) lion.

Members of the Felidae family are hypercarnivores that feed almost exclusively on flesh [9]. This is reflected in their shearing carnassial teeth [10], retractable claws, heightened sense of smell and vision, and optimized limb lengths [2, 11]. These characteristics are highly conserved across species of felids and are essential for the capturing and consumption of live prey [12]. Like other physical characteristics of felids, cat skulls share many features that are anatomically derived for predation. Wide zygomatic arches and post-orbital regions allow for strong jaw muscle attachment that confers high mechanical advantage and bite force greater than that of most other carnivores [2,9].

While the majority of predatory characteristics are conserved across cat species, there are distinctive differences between the mandibles of extinct sabretooth cats and extant conical toothed cats. The longer mandible found in sabretooth cats accommodates a larger gape for significantly larger canines [13-14]. There is considerable debate over how the mandible of sabretooth and conical tooth cats came to be so different, however, it has been suggested that sabretoothed and conical tooth cats were subjected to fundamentally different selective forces during their evolution [11].

Christiansen (2008) provided the first comprehensive analysis of lateral skull shape across both extant modern cats and extinct felids, including multiple species of sabretooth cats. Here, we build upon this work by characterizing skull shape in the dorsal and ventral regions of the skull, as well shape symmetry within the cranium. Geometric morphometric analysis was used to characterize skull shape change across 6 different species of felids, including *Smilodon*, NA lion, African lion, cheetah, leopard, and tiger. Our results indicated a significant shape difference in sabretooth cats and cheetahs, compared to other felid species. Notably, there was also evidence of asymmetry within the ventral cranium, specifically in the cheetah and NA Lion specimens, which may be attributed to a lack of genetic diversity in these species. Taken together, we provide evidence of cranial shape divergence across both extinct and extant members of Felidae, which likely evolved due to differences in predatory behaviors.

## Materials and Methods

### Specimen collection and origin

We characterized global shape and symmetry across six different species of Felidae. These species included the extinct *Smilodon fatalis* and *Panthera atrox*, as well as the extant *Acinonyx jubatus, Panthera leo, Panthera pardus, and Panthera tigris*. All Felidae skull specimens were obtained as photographs of museum specimens using a Nikon Coolpix 995 digital camera (Nikon Corporation, Tokyo, Japan). For the lateral mandible view, 7 cheetah, 7 leopard, 20 lion, 8 NA Lion, 4 NA Sabretooth, and 9 tiger specimens were available. For the lateral cranial view, 7 cheetah, 7 leopard, 20 lion, 9 NA Lion, 4 NA Sabretooth, and 9 tiger specimens were collected. For the dorsal cranial view, 7 cheetah, 7 leopard, 21 lion, 8 NA Lion, 3 NA Sabretooth, and 8 tiger specimens were collected. Finally, for the ventral cranial view, 3 cheetah, 3 leopard, 16 lion, 4 NA Lion, 3 NA Sabretooth, and 6 tiger specimens were collected. The *Smilodon fatalis and Panthera atrox* specimens were obtained from George Page Museum in Los Angeles, CA. All other extant specimen photographs were obtained from the Field Museum of Natural History in Chicago, IL.

### Landmark selection

Landmarks were selected on homologous cranial structures that could be reliably observed across all species of Felidae (Table 1). These landmarks were collected from the lateral mandible (7), lateral cranium (10), dorsal cranium (18), and ventral cranium (20) orientations. ImageJ software [15] was used to collect landmark data in two dimensions (2D; x and y coordinates) from all images.

**Table 1.**
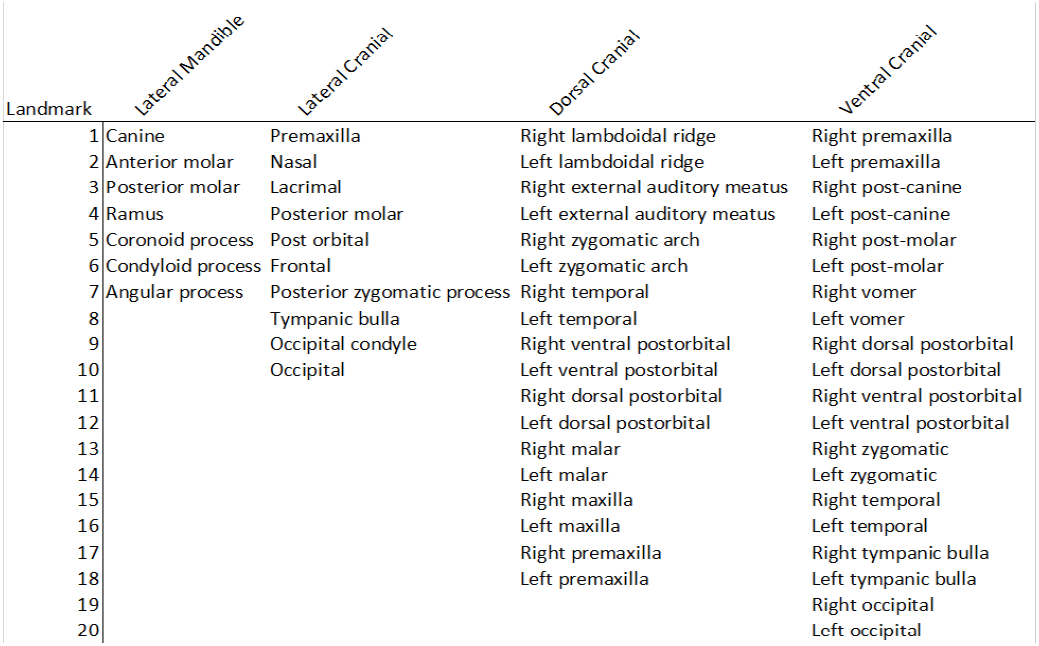
Homologous cranial landmarks selected in multiple orientations.

### Geometric morphometric analyses of skull shape

MorphoJ software [16] was used to conduct 2D geometric morphometric analysis of skull shape. A global analysis was performed for both the symmetric and asymmetric components of shape according to methods outlined in Powers *et al*. 2017. Briefly, a principal component analysis (PCA) was conducted to display shape variation across the various species of Felidae. Wireframe graphs of the principal component 1 (PC1) were generated to visualize the shape variation due to the principal component inclusive of the highest percentage of variation across species. Procrustes analysis of variance (ANOVA) was used to determine the relative amount of variation at each landmark (p<0.05).

## Results

### Divergence in lateral mandible shape is associated with the coronoid process

Landmark locations for all species were analyzed across four orientations of the skull. The Procrustes ANOVA for the lateral mandible indicated a significant difference in centroid size (p<0.0001) and shape (p<0.0001; Table 2) across different species of felid. Centroid size is a standard size variable that allows for a comparison of shape differences between specimens, regardless of differences in specimen size. Briefly, the centroid size can be found from the square root of the sum of squared distances of all landmarks from the center [17]. Principal component analysis (PCA) revealed 81% of this shape variation could be attributed to principal component (PC) 1 with a single landmark contributing to the majority of shape change (Figure 1A). We identified this landmark as landmark #5, which corresponded to the coronoid process (Table 1). This shape change was visualized using wireframe graphs (Figure 1A).

**Table 2.** Statistical analysis of shape change across Felidae species.

| **Procrustes ANOVA** |  |  |  |  |  |  |
| --- | --- | --- | --- | --- | --- | --- |
|  | *Effect* | *Sum of squares* | *Mean square* | *df* | *F* | *P* |
| **Lateral Mandible** | Individual | 1.10632281 | 0.01843871 | 60 | 51.98 | <0.0001 |
|  | Side |  |  |  |  |  |
|  | Ind*Side |  |  |  |  |  |
|  | Residual | 0.25895246 | 0.00035473 | 730 |  |  |
|  | *Effect* | *Sum of squares* | *Mean square* | *df* | *F* | *P* |
| **Lateral Cranium** | Individual | 0.23761825 | 0.00247519 | 96 | 7.16 | <0.0001 |
|  | Side |  |  |  |  |  |
|  | Ind*Side |  |  |  |  |  |
|  | Residual | 0.25986726 | 0.00034557 | 752 |  |  |
|  | *Effect* | *Sum of squares* | *Mean square* | *df* | *F* | *P* |
| **Dorsal Cranium** | Individual | 0.20446184 | 0.00159736 | 128 | 38.83 | <0.0001 |
|  | Side | 0.00187438 | 0.00011715 | 16 | 2.85 | 0.0005 |
|  | Ind*Side | 0.00526518 | 4.1134E-05 | 128 | 0.45 | 1.0000 |
|  | Residual | 0.14436841 | 9.2072E-05 | 1568 |  |  |
|  | *Effect* | *Sum of squares* | *Mean square* | *df* | *F* | *P* |
| **Ventral Cranium** | Individual | 0.08360334 | 0.00092893 | 90 | 28.59 | <0.0001 |
|  | Side | 0.00097897 | 5.4387E-05 | 18 | 1.67 | 0.059 |
|  | Ind*Side | 0.00292424 | 3.2492E-05 | 90 | 0.56 | 0.9995 |
|  | Residual | 0.06017291 | 5.7637E-05 | 1044 |  |  |

**Figure 1.**
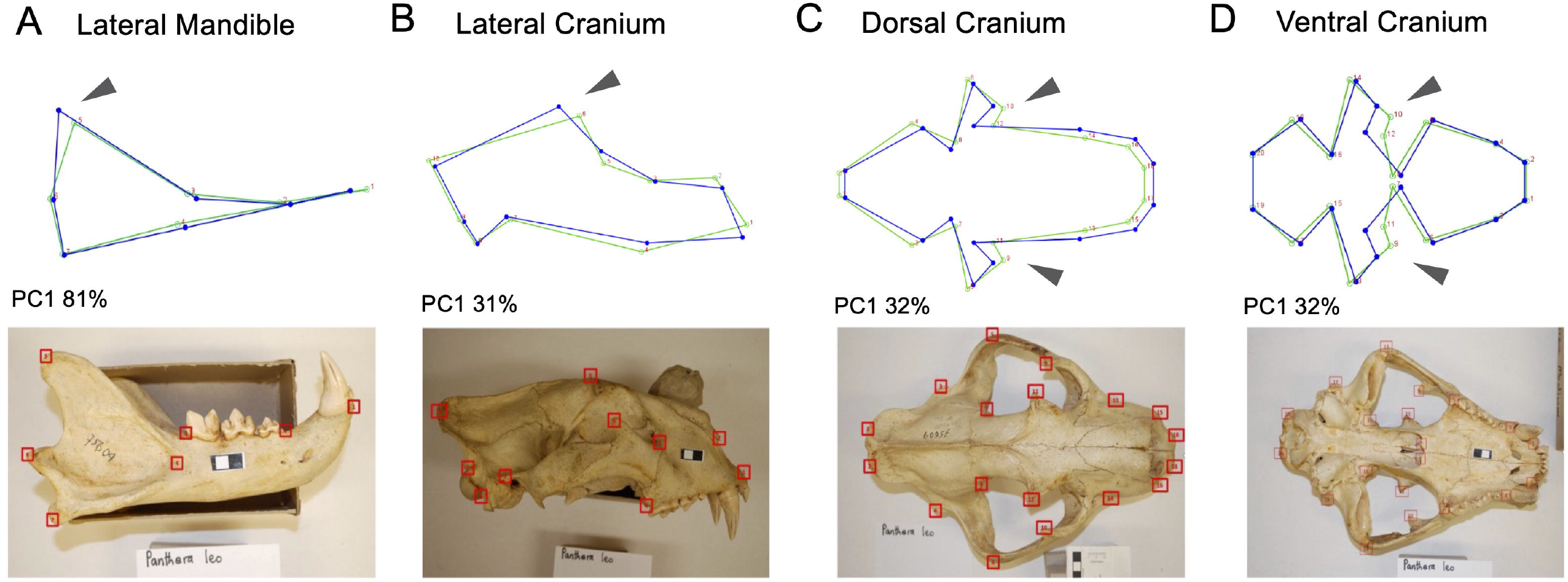
Wire frame graphs indicate regions of shape change in multiple skull orientations. Representative photographs from *Panthera leo* illustrate homologous landmarks (red) in each view (Table 1). Variation from a lateral mandible view at landmark #5, corresponding to the coronoid process indicates shortening of the mandible (PC1 81%, A). Variation from a lateral cranial view at landmarks #5 and #6 indicates a shortening of the postorbital and frontal bone (PC1 31%, B). Variation from a dorsal and ventral cranial view at landmarks #9-12 indicate shortening of the postorbital region (PC1 32%, C; PC1 32%, D).

### Two felids exhibit shortened postorbital and frontal regions

#### Lateral orientation

Procrustes ANOVA for the lateral cranium indicated a significant difference in centroid size (p=0.0146) and shape across the different species of felid (p<0.0001) (Table 2). An analysis of the principal component revealed that 31% of this variation could be attributed to PC1 (Figure 1B). Landmark #6, which corresponded to the frontal bone (Table 1), was identified as the region exhibiting the greatest shape divergence. The second greatest source of variation identified by the wireframe graph was landmark #5, which corresponded to the post orbital bone (Table 1). This landmark contributed to 21% of total shape variation (PC2). The change in shape at both landmarks was visualized using the wire frame graphs (Figure 1B).

#### Dorsal orientation

Procrustes ANOVA of the dorsal cranium landmarks indicated a significant change in skull size (p=0.0403) and shape (p<0.0001) (Table 2). Visualization of the dorsal cranium using wireframe graphs revealed 32% of total shape variation could be attributed to PC1 and landmarks #11 and #12 were pinpointed as major regions of shape change (Figure 1C). These landmarks were identified as the right and left dorsal postorbital processes (Table 1). Skull shape variation was also observed at landmarks #9 and #10 (Figure 1C). These landmarks accounted for 32% of total shape variation and were identified as the right and left ventral postorbital processes (PC2; Table 1).

#### Ventral orientation

Procrustes ANOVA of ventral cranium landmarks indicated an insignificant change in skull size (p=0.0974), however there was a significant change in shape (p<0.0001; Table 2). Visualization of the ventral cranium using wireframe graphs indicated 32% of total shape variation could be attributed to PC1, implicating landmarks #11 and #12 as sources of morphological change (Figure 1D). These landmarks were identified as the right and left ventral postorbital processes (Table 1). Skull shape variation was also visualized at landmarks #9 and #10 (Figure 1D). These landmarks accounted for 30% of total shape variation and were identified as the right and left dorsal postorbital processes (PC2; Table 1).

### PCA reveals shape differences in cheetah and NA Sabretooth specimens

PCA plots were utilized to identify species with the greatest source shape variation through ordination (Figure 2). Species plotted close together had greater shape similarity and species plotted farther apart had greater shape difference, with confidence ellipses overlayed (p<0.05). The PCA graph of the lateral mandible view revealed that the NA Sabretooth cluster was distinct from the other clusters of species (Figure 2A). Further, the PCA graph of the lateral cranial view revealed two clusters of specimens distinct from other species. These clusters corresponded to the NA Sabretooth and cheetah specimens. The PCA graph of the dorsal cranial view revealed a cluster of specimens distinct from other species that corresponded to the cheetah (Figure 2C). Finally, the PCA graph of the ventral cranial view revealed a cluster of specimens distinct from other species, corresponding to the NA Sabretooth (Figure 2D).

**Figure 2.**
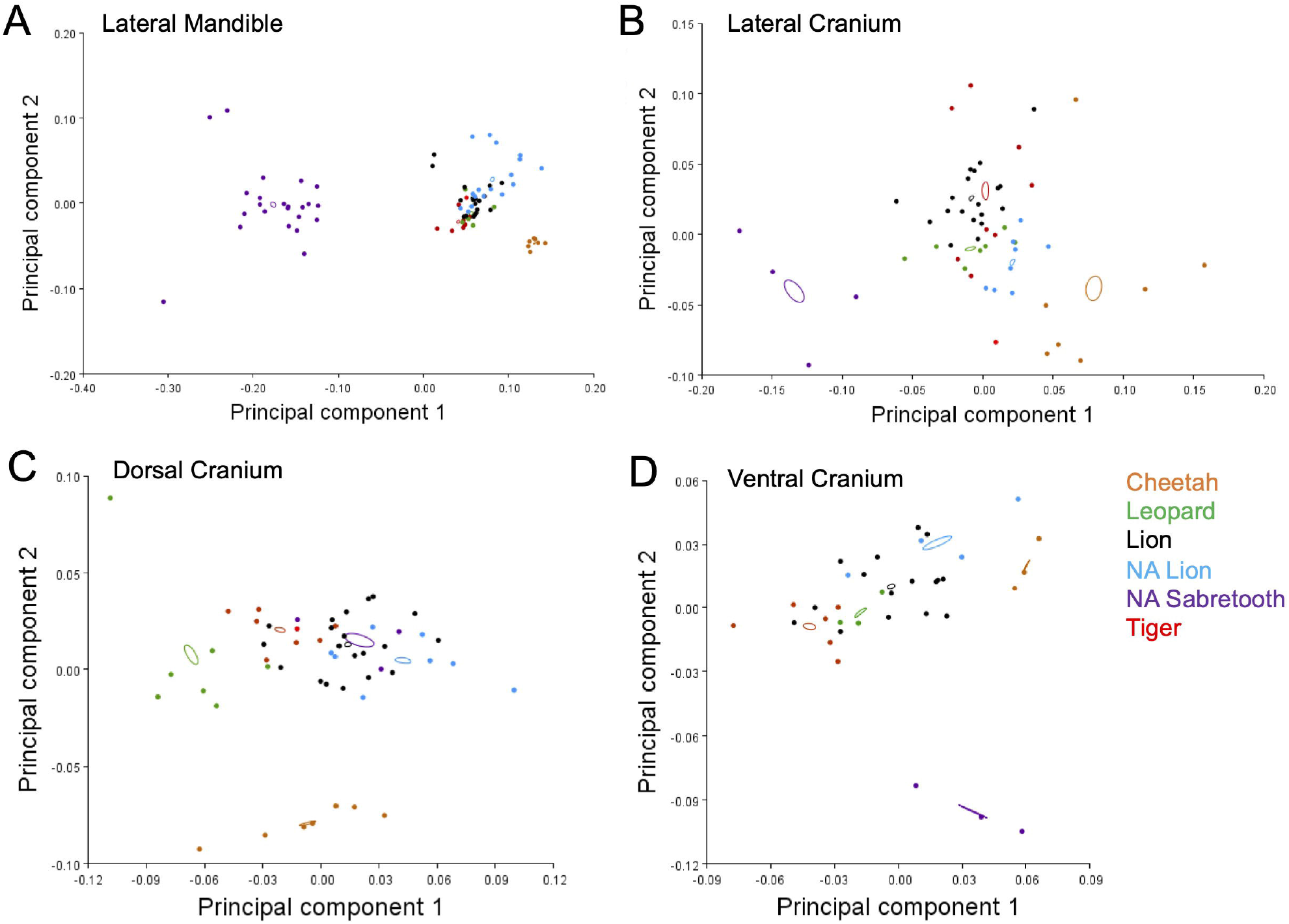
NA Sabretooth and Cheetah species exhibit the greatest shape variation across felids. PCA graphs of all four anatomical orientations indicate specimens with the greatest shape variation. The purple cluster of specimens in PC graph (Lateral mandible, A) indicate the coronoid process of the NA sabretooth is significantly shorter than other species of felid. The purple and orange cluster of specimens in PC graph (Lateral cranium, B) indicate a shortening of the postorbital region in the NA sabretooth and the cheetah. The orange cluster of specimens in PC graph (Doral cranium, C) and the purple cluster of specimens in PC graph (Ventral cranium, D) indicate a significant shape change in the postorbital region in both cheetahs and NA Sabretooth cats compared to other species of Felidae.

### Significant ventral asymmetry attributed to cheetah and NA Lion specimens

In addition to global shape change, skull shape symmetry was analyzed across multiple orientations of the cranium. A Multivariate Analysis of Variance (MANOVA) across all species was performed on the asymmetric components of shape [16]. The left and right sides in the lateral view could not be compared because only a single photo was taken for the lateral orientation. From the dorsal view, there was no indication of asymmetry in skull shape (p=0.3405). However, in the ventral view, we discovered a significant source of shape asymmetry (p=0.0002; Figure 3B). Further analysis of the asymmetrical component of the ventral cranium indicated asymmetry of the orbital region, primarily within the cheetah and NA lion species (Figure 3A).

**Figure 3.**
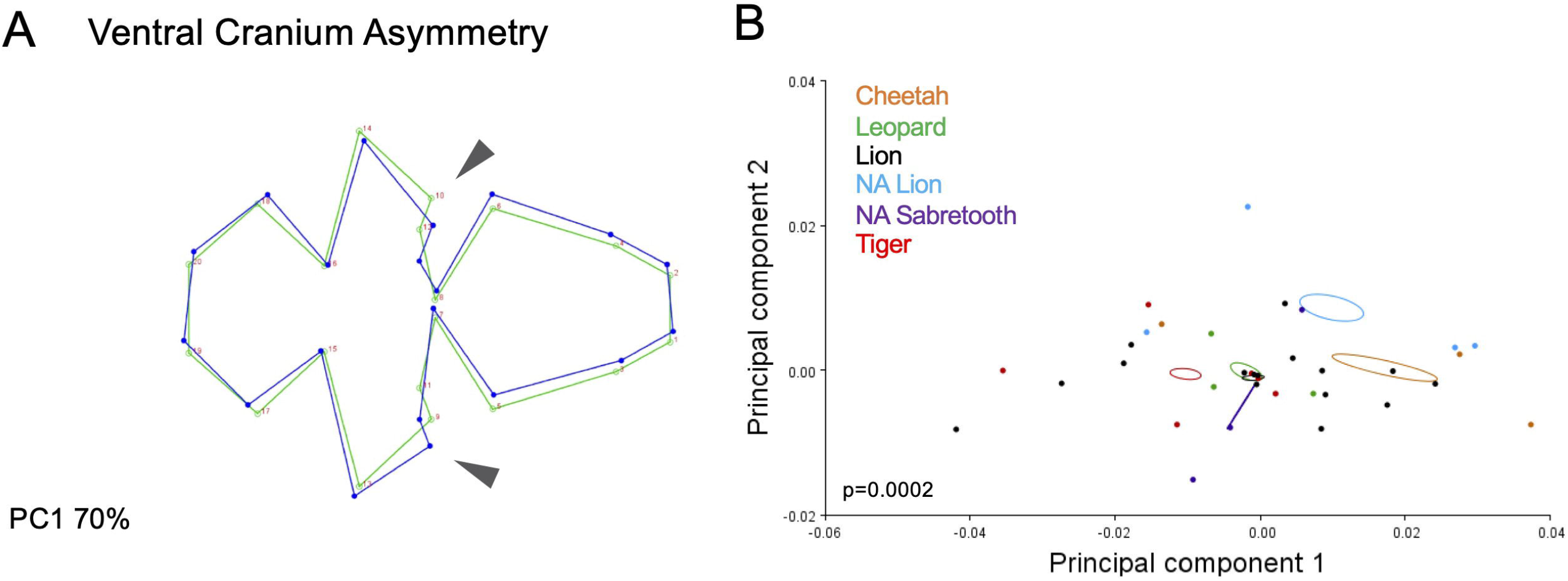
The NA Lion and Cheetah species exhibit significant cranial asymmetry. A wireframe graph of ventral asymmetry was constructed to visualize sources of significant asymmetry (A). The greatest sources of asymmetry can be visualized by landmarks #9-12. These landmarks correspond to the postorbital region of the skull (Table 1). A PCA graph of ventral asymmetry reveals significant asymmetry in two species of felid, NA Lion and Cheetah (B).

## Discussion

### Shortened coronoid process observed in the NA Sabretooth

Much of felid predatory morphology is conserved across species, including tooth and claw structure, as well as enhanced vision [9, 10, 12]. Our study utilized geometric morphometric analysis to identify changes in skull morphology across Felidae species. Procrustes and principal component analysis of the lateral mandible revealed a shortening of the coronoid process in NA Sabretooth specimens, compared to other felid species (Figures 1A; 2A). These results are consistent with other morphometric analysis of sabretooth morphology [11,18-19].

Due to the extinct status of NA Sabretooth cats, the predatory and feeding behaviors of this species are subject to speculation. Geometric morphometric analysis can be used to make inferences regarding the feeding behaviors in these predators. It has been proposed that the shortening of the coronoid process observed in NA Sabretooth mandibles enabled the large jaw gape necessary to accommodate their enlarged canines [18].

The shortening of the coronoid process in NA Sabretooth indicates a lower mechanical advantage and bite force, compared to extant felid species [18,20]. A longer coronoid process was observed in extant species of felids, which is associated with a high mechanical advantage and strong bite force necessary for occluding the throats of potential prey [19]. Another study of *Smilodon* morphology concluded that sabretooth skull shape was better suited for alternative methods of capturing prey. Their prey were likely subdued by powerful forelimbs and killed with a shearing bite from enlarged canines [14].

### Shape change in the postorbital region is attributed to NA sabretooth and cheetah

Analysis of the ventral cranium revealed significant shape change in the postorbital region toward the anterior skull (Figure 1D) is attributed to the NA Sabretooth (Figure 2D). The Felidae family generally have large orbits to assist in predation by enhancing vision [9]; however, most machairodont species, such as the NA Sabretooth, had smaller orbital regions and likely restricted their hunting to the daylight hours [1].

Analysis of the lateral and dorsal cranium revealed a shortening and enlargement of the orbital region of the cheetah skull compared to other species of felid (Figure 2B, C). This suggests a larger eye size and more anterior position on the skull. The change in eye size and position observed in the cheetah may be explained by their predatory behaviors. The cheetah is the fastest known land mammal on earth, reaching speeds of up to 90kmh [21-23]. Previous studies on visual acuity in mammals have found that increases in eye size are associated with increased visual acuity [24]. This increase in visual acuity would likely confer an advantage when tracking prey at high speeds. Despite their running prowess when reaching high speeds, the cheetah can only maintain maximum velocity for a short time, giving up after a few hundred meters [22]. Following the short exertion when hunting prey, the cheetah must rest, leaving them vulnerable to predation and kleptoparasites. To prevent predation and meal loss, some cheetahs are able to drag their prey to tall grasses [25-27]. Some cheetahs even utilize tall grasses to stalk prey before a hunt, which assists them by reducing run distances [25]. The anterior eye position observed in cheetahs may assist in navigating these grasses.

### Significant asymmetry identified in cheetah and NA lion specimens

A deeper analysis of skull symmetry from the ventral view revealed significant asymmetry, in the orbital region, that could be attributed to both the cheetah and the NA Lion (Figure 3A, B). Previous research examining skull symmetry in cheetahs found fluctuating levels of asymmetry, which were attributed to inbreeding and population bottleneck within the cheetah population [28]. An analysis of the complete cheetah genome confirmed a significant lack of genetic diversity within the species compared to other mammals [29]. Genetic analysis has also provided evidence for the existence of two possible bottleneck events within the evolutionary history of the cheetah [29]. Research has shown that population bottlenecks can result in developmental instability, increasing incidences of fluctuating asymmetry [30-31].

Our results also indicated skull asymmetry within the orbital region, of the NA Lion specimens. These results were unexpected. Not much research has been conducted examining the genetic makeup and skull symmetry of the NA Lion. It is worth noting that less ventral cranial specimens were available for analysis and discrepancies in shape symmetry may be due to smaller sample size. Future research into the genetic variation and skull symmetry of the NA Lion using a larger sample could provide an explanation for the asymmetry seen within our specimens.

## Acknowledgments

The authors are grateful for the support from the Xavier University Biology department, and in particular the summer research funding awarded to AN. This research was made possible by paleontological expeditions and access to museum specimens (George Page Museum in Los Angeles, CA and the Smithsonian Institution in Washington D.C.). AKP is supported by NIAMS (1F32AR076187-01).

## Notes

### Competing Interest Statement

The authors have declared no competing interest.

